# End-to-End deep structure generative model for protein design

**DOI:** 10.1101/2022.07.09.499440

**Authors:** Boqiao Lai, Matt McPartlon, Jinbo Xu

## Abstract

Designing protein with desirable structure and functional properties is the pinnacle of computational protein design with unlimited potentials in the scientific community from therapeutic development to combating the global climate crisis. However, designing protein macromolecules at scale remains challenging due to hard-to-realize structures and low sequence design success rate. Recently, many generative models are proposed for protein design but they come with many limitations. Here, we present a VAE-based universal protein structure generative model that can model proteins in a large fold space and generate high-quality realistic 3-dimensional protein structures. We illustrate how our model can enable robust and efficient protein design pipelines with generated conformational decoys that bridge the gap in designing structure conforming sequences. Specifically, sequences generated from our design pipeline outperform native fixed backbone design in 856 out of the 1,016 tested targets(84.3%) through AF2 validation. We also demonstrate our model’s design capability and structural pre-training potential by structurally inpainting the complementarity-determining regions(CDRs) in a set of monoclonal antibodies and achieving superior performance compared to existing methods.

## 1 Introduction

Computational protein design has been of great interest to the scientific community for decades. Designing protein macromolecules with specific function and structure is a highly sought after technique with broad application to therapeutics, biosensors, and enzyme engineering (Griss et al., 2014; Yang et al., 2021; Lu et al., 2022; Huang et al., 2016; Zhang et al., 2022a). However, despite years of effort and advancements, computational protein design still remains a very challenging problem.

Traditionally, the protein design pipeline is often regarded as a two step process where the practitioner first determines the protein backbone structure accommodating the specified structural and biochemical properties, and then designs the amino acid sequence with the given backbone structure. In this regime, the most successful applications rely heavily on template-based fragment sampling and domain-expert specified topologies for backbone determination and energy minimization based sequence design (Courbet et al., 2022; Cao et al., 2022). While this approach is widely adopted, there are apparent drawbacks. For example, the resulting backbone structure from the first step may not be optimal or designable and the designed sequence in the second step may not readily conform to the desired backbone template. As a result, the success rate for computational protein design remains relatively low (Huang et al., 2016; Anishchenko et al., 2021).

In recent years, progress in machine learning and deep learning research has contriburted to significant advances for protein modeling such as mutation effect estimation (Frazer et al., 2021; Meier et al., 2021; Riesselman et al., 2018), protein function prediction (Lai & Xu, 2021; Gligorijević et al., 2021), and structure prediction (Jumper et al., 2021; Baek et al., 2021; Wang et al., 2017).These advancements have also played a part in aspects of the computational protein design problem. For fixed backbone sequence design, a series of deep learning methods have emerged to improve conventional energy based approaches (Alford et al., 2017; Khatib et al., 2011) by directly incorporating structural information using SE(3)-equivaraint frameworks (McPartlon et al., 2022; Jing et al., 2020; Hsu et al., 2022). An array of recent works have studied the use of generative models for structure generation (Lin et al., 2021; Anand et al., 2019; Anand & Huang, 2018), however, these methods often generate topological constraints and rely on downstream tools for 3-dimensional structure determination. One of the major difficulties for direct coordinate generative modelling is properly accounting for the rotation and translation equivariance in the target conformation.

In the paper, we present a versatile VAE-based deep structural generative model that seeks to bridge the gap for robust computational protein design. Our method makes contributions to three fundemental aspects of protein design: First, we directly model protein structure in the 3-dimensional coordinate space which avoids downstream coordinate recovery in constraints based model. Second, Our method is universal such that it can model proteins of arbitrary size and thus exposes our model to the whole fold space while previous generative models are restricted to protein with certain size and can only be trained on a small subset of all available folds for a given model. Third, We address the translation and rotation equivariance in both the input space and the use of a locally aligned coordinate loss proposed by Jumper et al. (2021). It is important to note that structure entries in database such as the Protein Data Bank(PDB) (Berman et al., 2000) often represent a single sample from its conformational landscape. By conditioning on a given backbone structure, our model is able generate ***conformational decoys*** from the latent space. Combined with fixed-backbone sequence design models and accurate protein structure prediction tools, our model can enable efficient *in silico* design screening.

We evaluate our model on three fundamental design tasks. First, we demonstrate that our model can generate high-quality protein structures by comparing the generated 3-dimensional coordinates to experimentally determined structures. Next, we show that we can improve efficiency and success rate in conventional design pipelines by producing sequences recapitulating high-quality backbone templates derived by our model. Last, we corroborate our model’s design capabilities by inpainting the backbone structure of the complementarity-determining regions (CDRs) of monoclonal antibodies. For this task, we achieve state of the art results, and exemplify how our model can be used as a pre-trained structure encoder.

## 2 Related Work

### Deep Generative models

generative adversarial networks(GAN) (Goodfellow et al., 2014) and auto-encoding variational Bayes (VAE) (Kingma & Welling, 2013) are powerful generative frameworks that are used from image synthesis (Brock et al., 2018; Razavi et al., 2019) to language modeling (Bowman et al., 2015). Riesselman et al. (2018); Shin et al. (2021); Frazer et al. (2021) also used VAE for protein sequence design and variant effect prediction. Most of the aforementioned structure generative models also used VAE or GAN as their generative framework.

Our model is based on the VAE framework where we first encode the invariant protein structure representations(Methods) into the latent space then a decoder is used to generate the corresponding 3-dimensional coordinates from the latent space. This approach allows our model to generate flexible protein conformations conditioned on the input backbone constraints. Trained on masked input constraints, our model can easily be adopted for backbone inpainting for structure design.

### Protein structure generative models

Existing generative methods for protein structure design focus on generating invariant topological constraints such as inter-residue contacts and distance maps which are then converted into three dimensional structures via some downstream tools. (Lin et al., 2021; Anand et al., 2019; Anand & Huang, 2018; Karimi et al., 2020; Huang et al., 2021; Guo et al., 2021). Though these methods have garnered some success, there is no a priori guarantee than the generated constraints are geometrically viable. Consequently, the resulting three dimensional structures are often of low quality or biochemically infeasible. Moreover, it is generally not possible to perform conditioned structure generation on these models.

While Eguchi et al. (2020) proposed the first direct coordinate structure generative model, its application is limited to proteins of a fixed length, and is trained only to recover inter-atom distance and torsion maps. In contrast, our model, addresses rotation and translation equivariance in both the input and output space by distilling invariant representations of protein geometry and by using a locally aligned coordinate loss function to perform gradient optimization directly on the coordinate space. In this way, our model can directly and flexibly model the 3-dimensional structure and generate conformational decoys.

### Fixed backbone sequence design

For designing an amino acid sequence with a given backbone structure, it is conventional to use energy based optimization schemes (Alford et al., 2017; Khatib et al., 2011). This approach comes with several drawbacks, such as the exponential search space and limited capability to model higher order interactions. Recently, many deep learning based models have explored the use of rotation equivariant frameworks (McPartlon et al., 2022; Jing et al., 2020; Hsu et al., 2022) and invaraint 3-dimensional voxel-based approaches (Qi & Zhang, 2020; Anand et al., 2022). Although our model does not directly design the sequence alongside the conformation decoys, we show that sequences designed from the resulting templates can be more reliable than those designed from a single backbone structure.

## 3 Results

### 3.1 Overview

In this section we illustrate the overall model design as shown in Figure 1. To prepare for input features, each protein structure is distilled into invariant pairwise representations of inter-residue distance and orientations as described in (Yang et al., 2020) and scalar representations of amino acid sequence and backbone torsion angles. This input is then fed through an encoder network which produces a latent representation of each residue. These representations are reassembled and passed to a decoder module which reconstructs the backbone coordinates. The process is described in full detail in Method Section 5.

**Figure 1:**
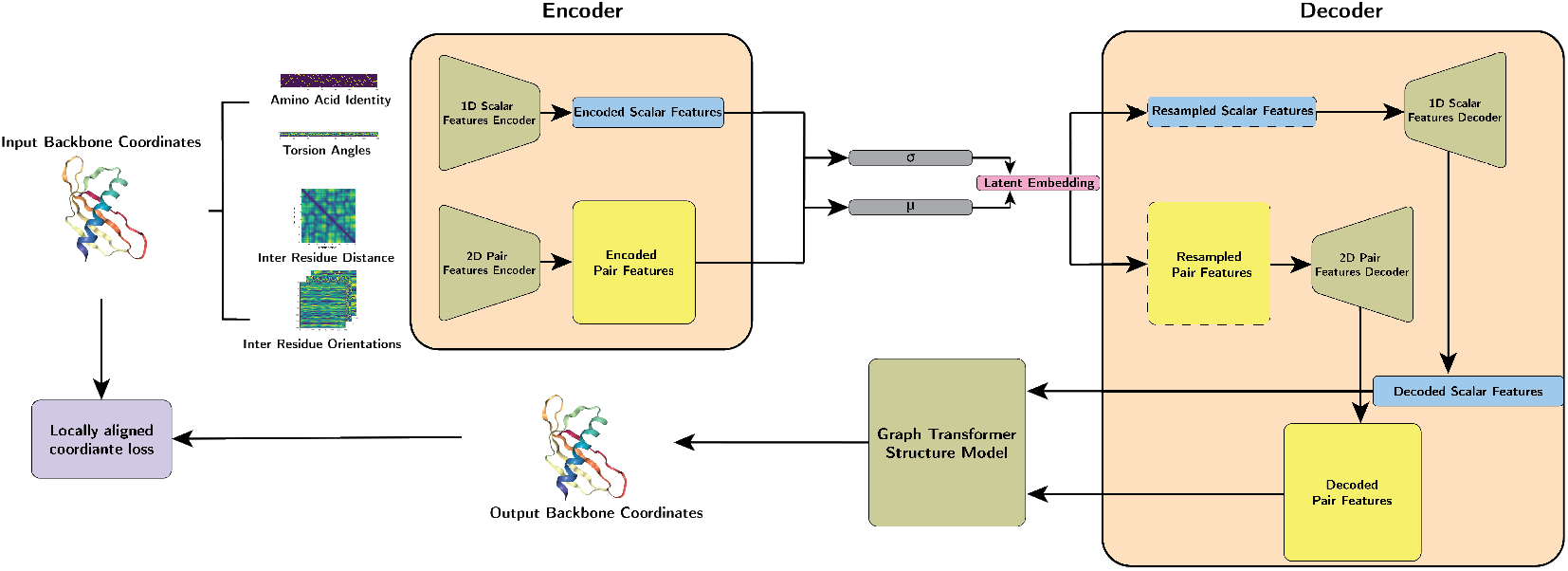
Model Overview. Structures are first distilled into pairwise (inter-residue distance & orientation) and scalar (torsion angles & amino acid sequence) features, Then our model separately encodes the pairwise and scalar features into the latent space. The latent representation is then sampled and decoded by the pair and scalar feature decoder before being fed into a graph-transformer based structure module for coordinate generation. A locally aligned loss is then computed between the reconstructed coordinates and the native input structure.

We first demonstrate our model’s ability to generate realistic backbones structures through comparing with a set of experimentally determined and structurally independent structures. Then we show how the conformational decoys generated by our mode outperforms native structure backbones for designing structure confining amino acid sequences. Lastly, we tested out model’s structure design capability by testing CDR inpainting on a set of monoclonal antibodies.

### 3.2 End-to-End structure reconstruction & generation

In this section, we will demonstrate our model’s ability to reconstruct high-quality protein 3-dimensional structures from invariant protein representations. The most crucial capabilities for structure generative models is to construct topologically feasible protein conformations such that the generated backbone structures are viable candidates for downstream design pipelines. For this propose, we presented structural similarity metrics including the local distance difference test (lddt) score (Mariani et al., 2013), TM-score (Zhang & Skolnick, 2004), root mean square deviation (RMSD), and the *L*_1_ norm of inter-residue distance between the reconstructed structures and the native input structures to measure how well our models reconstruct the input structures.

Our model is able to consistently construct high-quality structure coordinates of the target proteins up to *L* = 500 residues as shown in Figure 2(A & B). To further inspect the geometric feasibility of the reconstructed proteins, we examine the torsion angle distributions and compared it to the native structures as shown in Figure 2(C) where we notice the torsion angles from the generated structures are mostly in the feasible regions and matches up closely to the native torsion angels. To illustrate how our model’s learned latent representation of the input protein contains useful information in its fold space, we plot the t-SNE embedded components colored by its CATH class annotations in Figure 2(D). Structures that share the same CATH fold class are clustered in the latent space accordingly.

**Figure 2:**
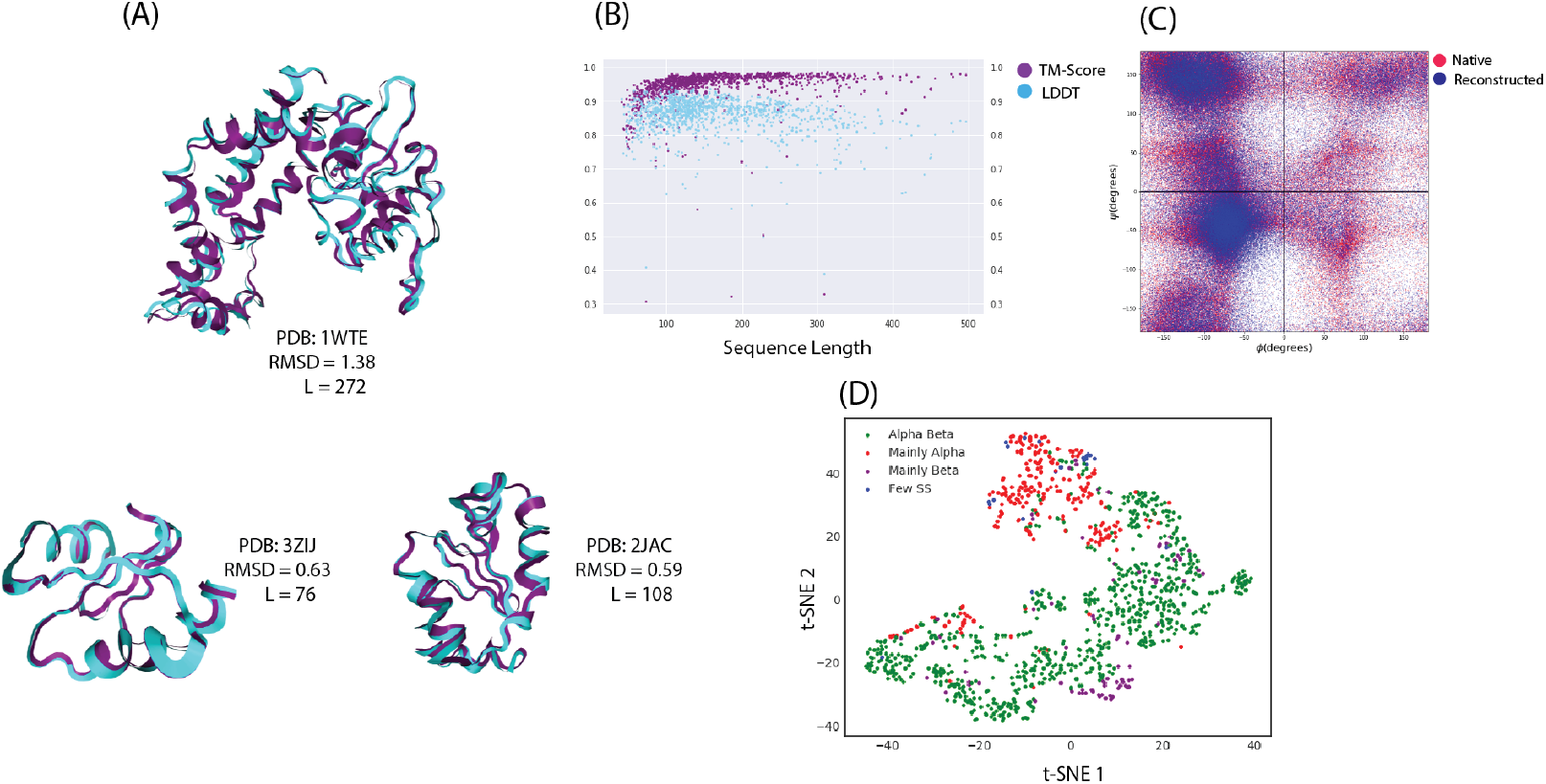
(A) Examples of reconstructed backbone structures of different folds and sizes. 1WTE (top), 3ZIJ (bottom left), 2JAC (bottom right). (B) LDDT (blue) and TM-score (purple) against native structure vs. the length of the test target. (C) Ramachandran plot of the native (red) and reconstructed(blue) structures. (D) t-SNE embedding of the latent space for the test targets colored by their CATH class annotation.

To understand how different input features impact model performance, we compared our model with different features ablated in table 1. When both inter-residue distance and orientation are used along with the amino acid sequence, our model achieved average RMSD of 1.122 and TM-score of 0.941 at experimentally comparable resolution. We notice model performance degrades when we remove the inter-residue orientation as input features. This suggests our model relies on spatial orientation information to accurately construct the structure coordinates. It is worth noting that even tough our model achieves the best performance with amino acid sequence input, our model has comparable result when operate under sequence free mode for backbone-only decoy generation. In addition, we also compared the short, medium, long range contact accuracy where the predicted contacts are derived from the inter-residue distance of the generated 3-dimensional coordinates as shown in table 3.

**Table 1:** Average structural similarity metrics evaluated on the test targets over different models

| Model | Iddt↑ | TM↑ | RMSD↓ | Distance LI↓ |
| --- | --- | --- | --- | --- |
| CoordVAE CNN | 0.472 | 0.420 | 9.049 | 2.666 |
| CoordVAE w/o orientation | 0.604 | 0.649 | 4.429 | 1.413 |
| CoordVAE w/o seq | 0.788 | 0.905 | 1.489 | 0.699 |
| CoordVAE | **0.841** | **0.941** | **1.122** | **0.502** |

### 3.3 Conformational decoy sampling for robust CPD

In this section, we describe how our structural generative model can be used for robust *de novo* protein design. The design pipeline using conformational decoys is outlined in Figure 3. A primary design target backbone structure is provided as the design template, such a template can be generated by topology design program such as TopoBuilder (Sesterhenn et al., 2020) or other template based methods. Our structure generative model(CoordVAE) will embed the input backbone into a latent representation then multiple conformational decoys are generated by the structure decoder to form a library. A sequence library is then produced by running a fixed backbone inverse folding program on the decoy library. Each sequence will then be folded and validated by a structure prediction oracle such as AlphaFold (Jumper et al., 2021), and those which pass the structure validation criteria will be used to form a structure validated sequence library for further filtering and downstream experimental screening. Our structure model can generate thousands of decoys per minute on a single GPU for fast and efficient library curation.

**Figure 3:**
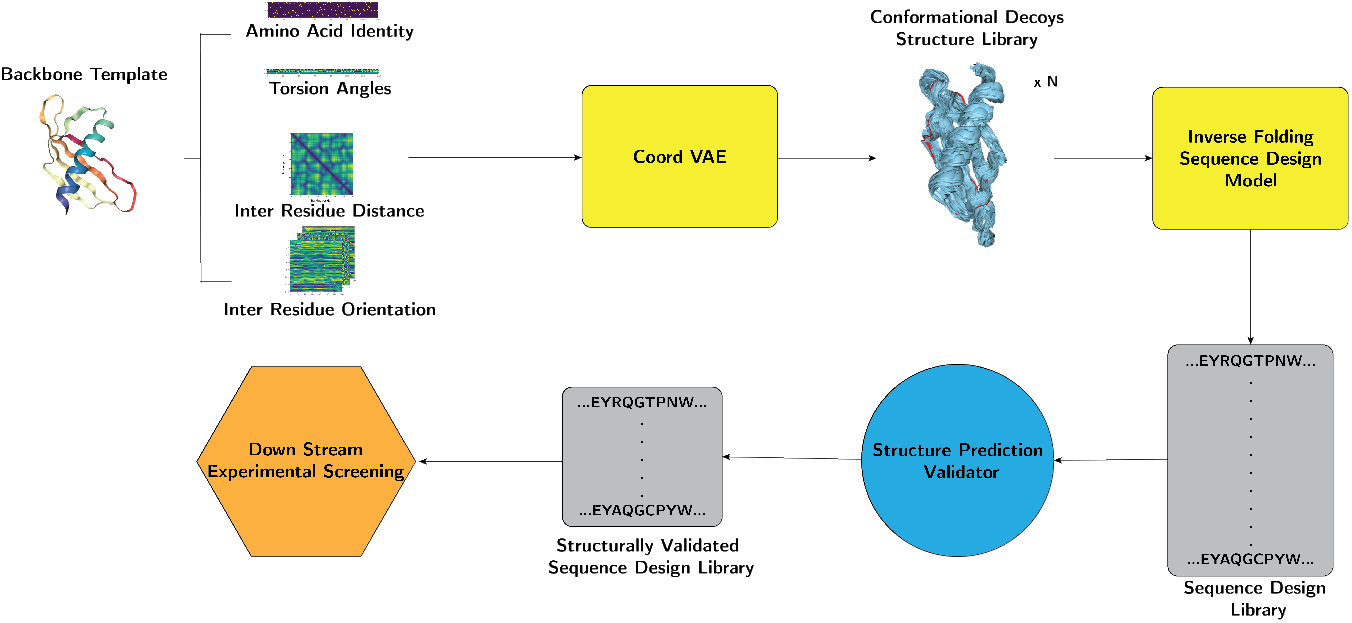
Outline of robust protein design framework using conformational decoy library. Starting with a primary backbone structure target, our structure generative model embeds it into a latent space and decodes into conformational decoys to form a structure library. Using a fixed backbone sequence design program on the backbone structure library, one can obtain a preliminary sequence library. To filter and structurally validate the primary sequence library, a structure prediction oracle is used and the validated sequence library is ready for further downstream tasks.

To elucidate how sequences designed from the conformational decoys compare to sequence designed from native backbones, we folded both sets of sequences with AlphaFold2 (Jumper et al., 2021). We provide examples in Figure 4(A) where our designed sequence folded more successfully than the native sequence when MAS and template information is not available. Particularly, as shown for PDB:3G67, the native sequence is folded with RMSD = 36.26 while our best decoy designed sequence folded with RMSD = 2.56 with high confidence. We observed that our model can consistently improve designed quality regardless of the size of the target backbone. In Figure 4 (B), we deployed the design pipeline to a set of 211 backbone targets and plotted the TM-score of the predicted structure compared to native sequence(bottom), and sequence designed from the native backbone structure(top). Sequences designed from our pipeline outperform sequences designed from fixed native backbones in 856 out of the 1,016 tested targets(84%). To confirm that our structure generative model does not simply generate noisy versions of the input structure, we compared with sequences designed from backbone coordinates with added random Gaussian noise. As shown in Figure 6(B), the results for conformatioal decoys are clearly favorable to those of noisy backbones. Additional results and examples can be found in the appendix Fig S1-S3.

**Figure 4:**
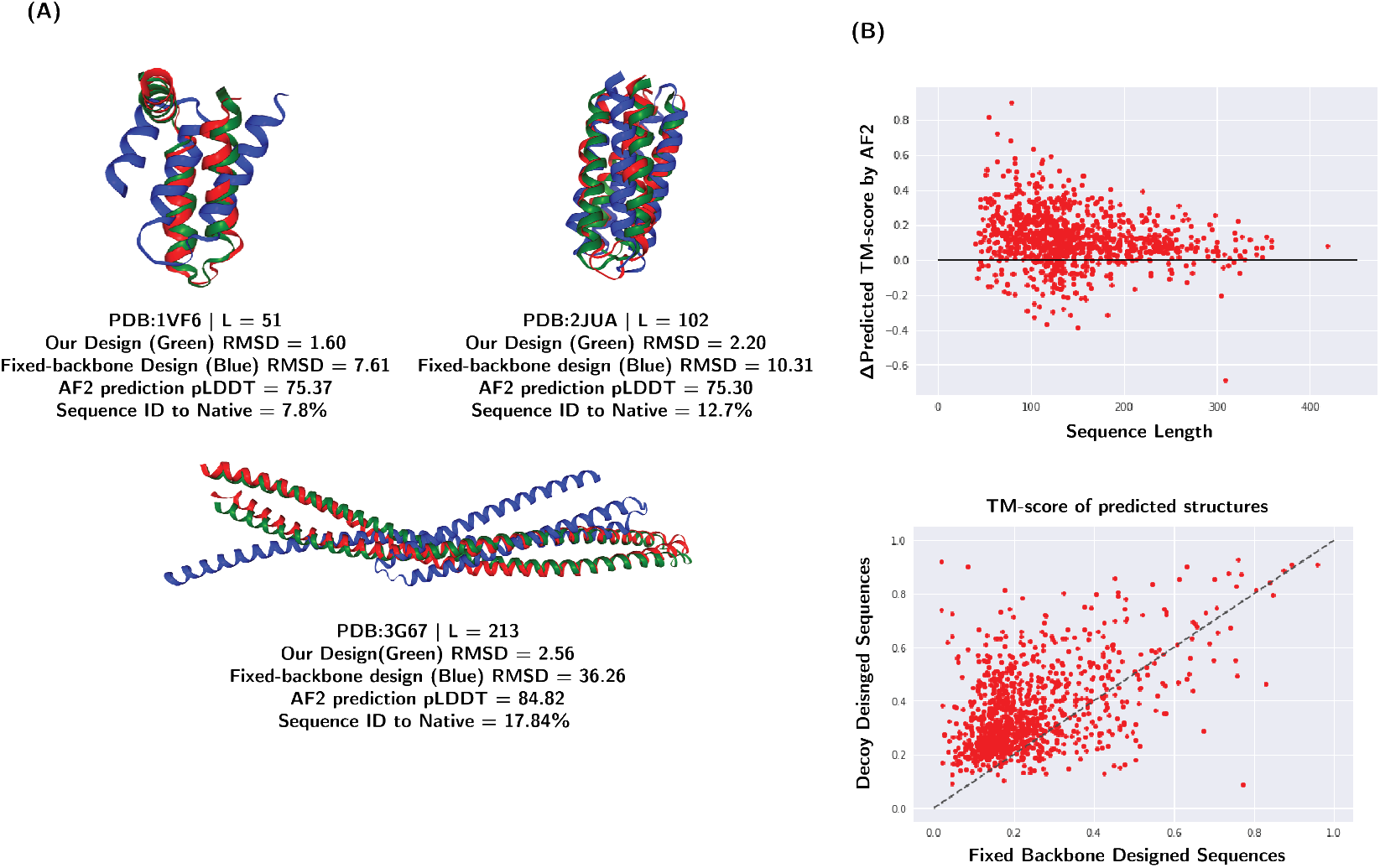
Examples of protein designed from conformational decoys. **(A)** AF2 folded sequence designed from conformational decoys. Overlay of AF2 predicted structure of decoy designed sequence & native backbone designed sequence (green & blue), native backbone(red) for PDB:1VF6 (top left), PDB:2JUA (top right), PDB:3G67 (bottom). **(B)** Scatter plots of TM-scores between the best folded conformational decoy designed sequences and sequences designed from the native backbones(bottom). ∆ TM-score between decoy designed sequences and fixed backbone designed sequences versus the target size.(top)

**Figure 5:**
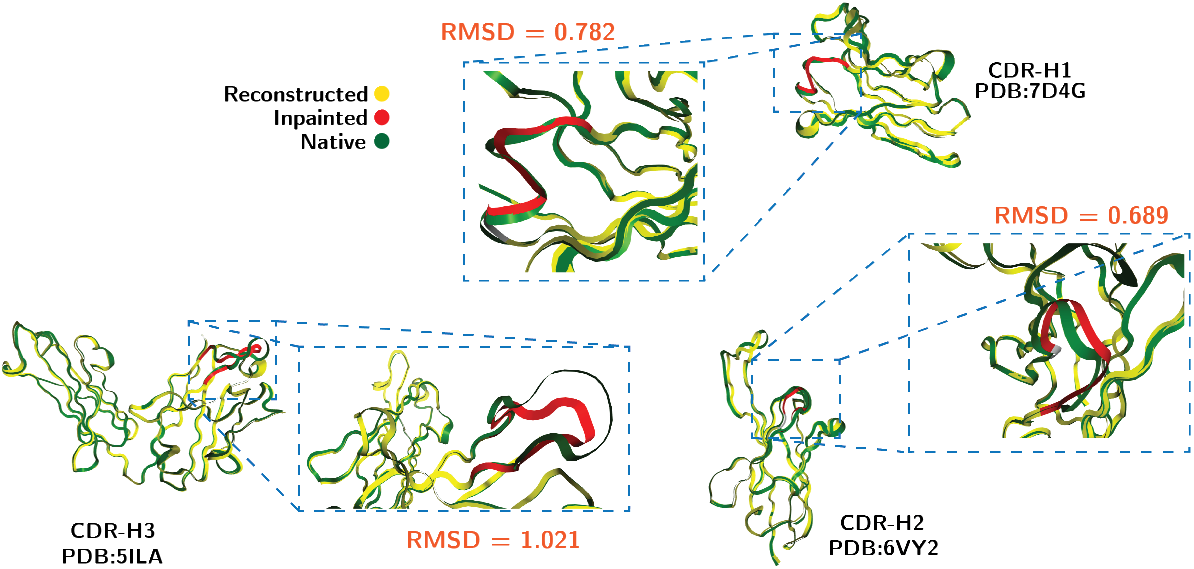
Examples of antibody structure inpainting: overlay of the CDR-H3 region of PDB:5ILA (bottom left) native structure(green), unmasked region (yellow) and masked inpainting region (red) reconstruction with RMSD = 1.021. overlay of the CDR-H2 region of PDB:6VY2 (bottom right) native structure(green), unmasked region (yellow) and masked inpainting region (red) reconstruction with RMSD = 0.689. overlay of the CDR-H1 region of PDB:7D4G (top) native structure(green), unmasked region (yellow) and masked inpainting region (red) reconstruction with RMSD = 0.782

**Figure 6:**
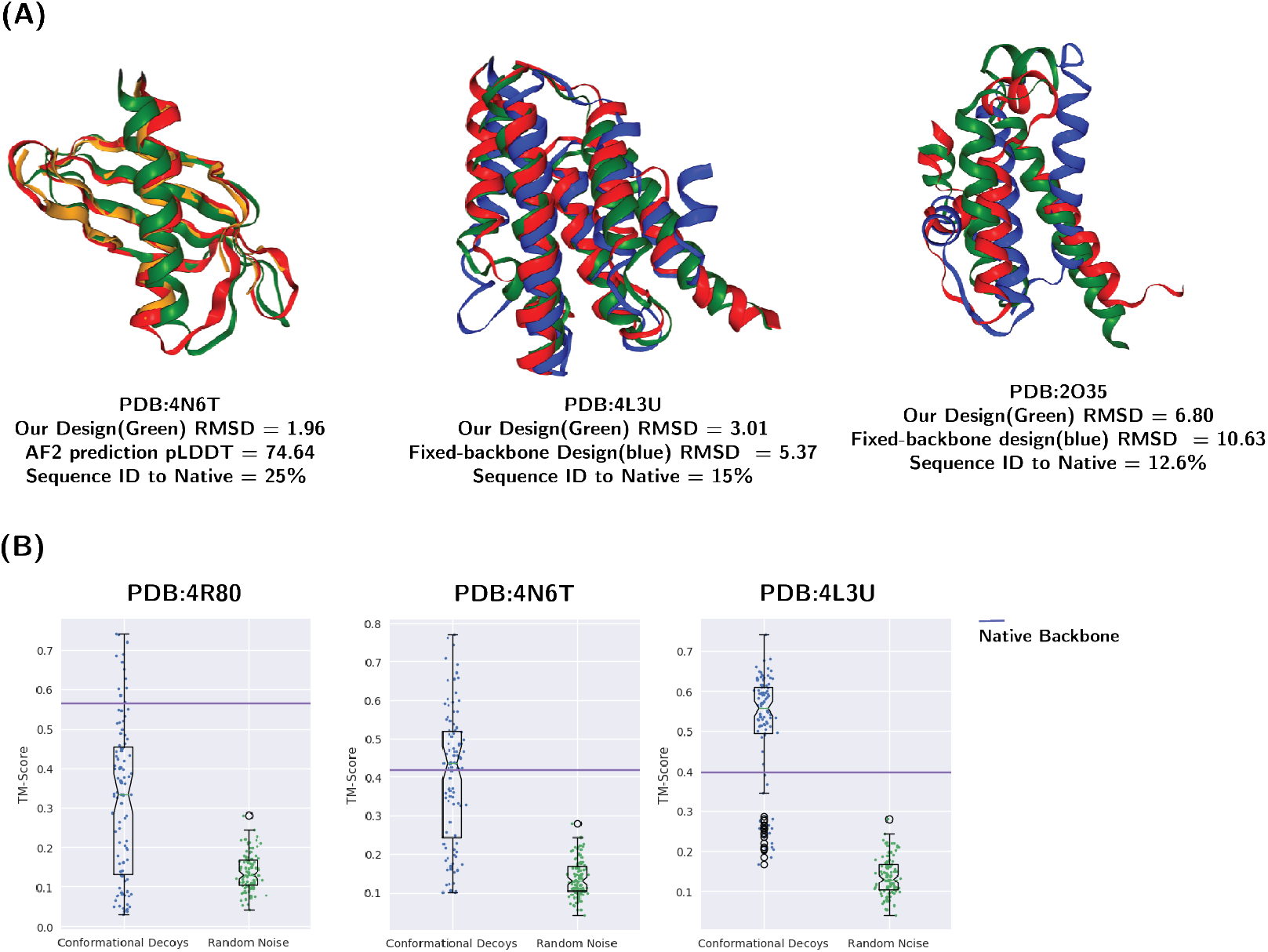
**(A)** Examples of AF2 predicted designed sequences and native sequences. **(B)** TM-score distributions of decoy designed sequences vs. noisy backbone designed sequences

In summary, we show that conformational decoys generated by our model can be used for robust protein design by providing a structure library for downstream fixed-backbone sequence design applications. Our experiments, show that structure conforming sequences can be reliably designed from the conformational decoys compared to single backbone targets. With this approach, we hope to present a new path towards robust and efficient protein design.

### 3.4 Structure inpainting for antibody design

Monoclonal antibodies are important targets for therapeutics development and considerable effort has been dedicated by the community towards computational antibody design (Pinto et al., 2020; Jin et al., 2021; Shin et al., 2021; Adolf-Bryfogle et al., 2018). While previous methods mostly focused on sequence design, we adopt our structure generative models to inpaint the complementarity-determining regions (CDRs) for structure based antibody design. In this section we show how our model can be used for structure design with a hybrid mask scheme and provide a direct comparison with existing structure design methods.

To inpaint protein structures, we employed three types of masks; linear, spatial and random and a hybrid masking strategy(see details in Section 5.1) that impose masks in the input scalar and pair representation and the completed structure will be recovered by the structure generative model. To further test the ability of our model to distill meaningful representation from a larger fold space, we pre-traiend our model on the CATH4.2 dataset and fine-tune the model with a set of monoclonal antibody structures.

We first tested our model’s inpainting performance on three CDRs(H1, H2, H3) with models trained on the CATH4.2 dataset without fine-tuning on antibody structures. Our best model trained on hybrid masking scheme achieved RMSD of 2.903, 3.223, and 3.889 on CDR-H1, CDR-H2, and CDR-H3 respectively as shown on table 2. Next, we test models trained on antibody structure with masked CDRs, without CATH4.2 pre-training, our model achieved RMSD of 1.55, 1.00, 1.55 respectively. With model pre-trained on the CATH4.2 dataset and fine-tuned on antibody structure with masked CDRs, our model achieved RMSD of 0.81, 0.85, 1.35 compare to state-of-the-art method RefineGNN (Jin et al., 2021) with RMSD of 1.18, 0.87, 2.50 respectively. To avoid information leakage, we filtered the CATH4.2 dataset with the antibody test sets for redundancy in pre-training.

**Table 2:** Structure inpainting performance on the test monoclonal antibody dataset in CDR-H1(left), CDR-H2(middle), CDR-H3(right) across different models.

| Model | Training Data | CDR-H1 |  | CDR-H2 |  | CDR-H3 |  |
| --- | --- | --- | --- | --- | --- | --- | --- |
| | | liddt↑ | RMSD↓ | liddt↑ | RMSD↓ | liddt↑ | RMSD↓ |
| CoordVAE(10AA linear) | CATH4.2 | 0.587 | 2.847 | 0.588 | 2.864 | 0.498 | 4.427 |
| CoordVAE(spatial) |  | 0.515 | 3.944 | 0.514 | 3.914 | 0.487 | 4.350 |
| CoordVAE(Random) |  | 0.447 | 4.216 | 0.534 | 4.152 | 0.450 | 5.185 |
| CoordVAE(mixed mask) |  | 0.591 | 2.903 | 0.575 | 3.223 | 0.530 | 3.889 |
| CoordVAE | SABDAB | 0.81 | 1.55 | 0.872 | 1.00 | 0.809 | 1.55 |
| **CoordVAE(Transfer)** |  | **0.88** | **0.81** | **0.90** | **0.85** | **0.823** | **1.35** |
| AR-GNN |  | N/A | 2.97 | N/A | 2.27 | N/A | 3.63 |
| RefineGNN |  | N/A | 1.18 | N/A | 0.87 | N/A | 2.50 |

**Table 3:** Average short(S), medium(M), long(L) contact accuracy evaluated on the 1,120 test targets over different models

| Model | Contact(S)↑ | Contact(M)↑ | Contact(L)↓ |
| --- | --- | --- | --- |
| CNN Baseline | 0.321 | 0.35 | 0.193 |
| CoordVAE w/o orientation | 0.500 | 0.436 | 0.378 |
| CoodVAE w/o seq | 0.875 | 0.833 | 0.785 |
| CoordVAE | **0.885** | **0.857** | **0.819** |

To summarize, our structure generative model can be adopted to perform structure inpainting by properly masking the input features. Through our experiment, we show the hybrid masking scheme can improve design performance and our model can adequately inpaint antibody CDRs. With pre-training on large structure databases, our model outperforms state-of-the-art antibody structure design models and we observe that our model has the largest improvement on longer inpainting regions(CDR-H3).

## 4 Discussion

In this paper, we presented a versatile VAE-based deep protein generative model for generating 3-dimensional protein structures. We demonstrated our model’s ability to construct realistic and viable designs from invariant input representations, which overcomes many limitations in existing protein structure generative models. Specifically, our model can directly generate protein structure in 3-dimensional coordinate space, accounting for rotation and translation equivariance while most existing models generate only topological constraints and rely on downstream tools for coordinate recovery. Using this to our advantage, we showcased how the generated conformational decoys can be used for robust and efficient computational protein design. Moreover, our method offers an tenable framework for latent structure representation and structure-based pre-training. We leverage our method’s ability to flexibly model proteins of arbitrary size by experimenting with pre-training for structure design and saw promising results. Last, we described how our model can be adopted for structure inpainting and design with a hybrid masking scheme. Tested on a set of monoclonal antibody structures, we saw encouraging results on structure based antibody design.

As a versatile structure generative model, we see many potential applications based on our framework. First, structure-based pre-training has recently been explored in graph-based models (Zhang et al., 2022b) with invariant representation and objectives, our model can be adopted for equivariant geometric protein representation learning with explicit 3-dimensional geometric information. We preliminary tested this idea in our inpainting experiment and result suggest such pre-training can improve downstream task performance. Due to limited compute and other constraints, our experiment only uses the CATH4.2 dataset for pre-training purposes and it will be interesting to explore pre-training with extensive protein structure databases such as the AFDB (Varadi et al., 2022) in other protein design and modeling settings. Second, it is relatively easy to extend our model to model protein complexes for protein interface design and flexible protein-protein docking.

Despite the promising results we presented in this study, we are aware that there are certain limitations to our structure generative framework. For example, our model cannot perform context free structure generation despite training with large number of protein folds, this hinders our model’s ability to generate completely novel protein folds without backbone conditioning.

## 5 Methods

### 5.1 Protein representations

We represent a protein as a complete molecular graph 𝒢 = (𝒱, ℰ) where 𝒱 consists of scalar residue features *v*_*i*_ and ℰ consists of pairwise features *e*_*ij*_ between residues *i* and *j*.

#### Scalar Features

The nodes *v*_*i*_ of our input graph correspond to protein residues *i* ∈ {1…*n*}. The input scalar feature 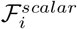 associated with residue *i* consists of amino acid identity and dihedral angle encodings:

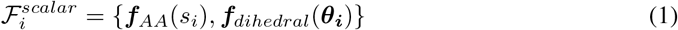

The first, ***f***_*AA*_ (*s*) ∈ ℝ^20^, is a one-hot encoding of the amino acid type *s* using 20 bins for each naturally occurring amino acid. The second, ***f***_*dihedral*_(***θ***) ∈ ℝ^6^, is an encoding of the three dihedral angels with Fourier features {sin, cos} ◦ {*ϕ*_*i*_, *ψ*_*i*_, *ω*_*i*_}, where *ϕ*_*i*_, *ψ*_*i*_, *ω*_*i*_ are dihedral angles computed from the coordinates from *C*_*i−*1_, *N*_*i*_, *C*_*αi*_, *C*_*i*_, *N*_*i*+1_ atoms.

#### Pairwise Features

for a given pair of residues i and j, we define the edge e_*ij*_ features as

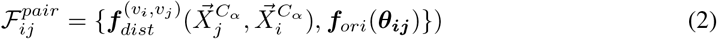

The first encoding, 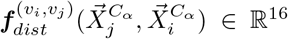 is the distance encoding that embeds the interresidue distance 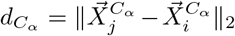 with 16 Gaussian radial basis functions with centers evenly spaced in [0, 20]A° as described in (Jing et al., 2020). ℱ_*ori*_(*θ*) ∈ ℝ^3^ is the encoding of the angle *θ* performed in the same manner as the backbone dihedral encoding for residue features. The input angles *θ*_*ij*_ *∈* { *ϕ*_*ij*_, *ψ*_*ij*_, *ω*_*ij*_ } are pairwise inter-residue orientations defined in (Yang et al., 2020). To produce pairwise orientation information, we impute a unit vector in the direction *Cβ*_*i*_ − *Cα*_*i*_ before computing the respective angles. The imputed vector is calculated as in (Jing et al., 2020) using

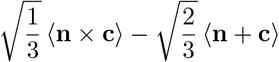

where ⟨*x*⟩ = *x*/∥*x*∥_2_, **n** = *N*_*i*_ − *Cα*_*i*_, and **c** = C_*i*_ − *Cα*_*i*_.

#### Masking Schemes for structure inpainting and pre-training

To help facilitate our model’s ability to inpaint protein structural regions, we employ a masking scheme for a given protein denoted as *Mask(n)*. We implemented three types of masks:

**linear** mask where a random residue is selected uniformly at random with *p* = 1/*L* where *L* is the length of the respective protein and the mask will then span 10 residues around the selected one. **spatial** mask where a random residue is selected uniformly at random with *p* = 1/*L* and all residues within 12A° are masked.

**random** mask where each residue will be masked at the rate of *p* ∼ *Uniform*(0, 0.5) where p is sampled for each individual residue.

For each masked residue, we use zero masks in both the scalar and pairwise features such that 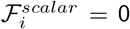 and 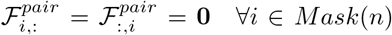. In the hybrid masking training strategy, 50% of the input proteins are mask with one of the aforementioned masks with equal probability. We also analyzed the effect of various masking probability and strategies, see the Appendix for more details.

### 5.2 Model Architecture

#### Feature encoders & decoder

The CoordVAE model uses different architectures for its encoder and decoder networks. The encoder network consists of a 1D feature module for the scalar input and a 2D feature module for the pairwise input using dilated convolution network architecture (Yu et al., 2017).

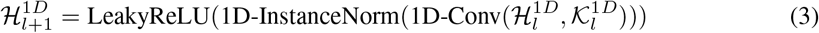

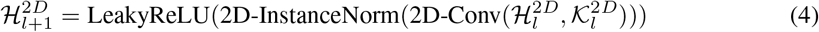

Where 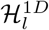 and 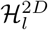 denotes the scalar and pairwise representation at layer l respectively and 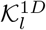 and 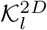 denotes the kernel size for the scalar and and pairwise convolution. Each convolutional block applies a leakyReLU nonlinearity with negative slope parameter set to 0.2, and instance normalization (Ulyanov et al., 2016).

The decoder network applies a mirrored architecture to the latent scalar and pairwise representations. Finally, the decoded scalar and pairwise features are processed jointly with a graph transformer described in (Shi et al., 2020) for coordinate generation.

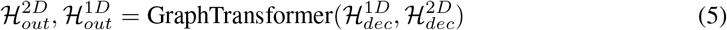

Where 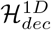 and 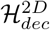 are the scalar and pairwise output of the decoder network respectively. For Coordinate generation, we process the output of the graph transformer module with a dense projection.

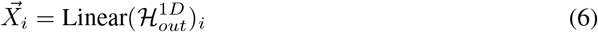

Where 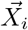 are the backbone coordinates of residue i and the projection is implemented with as a dense layer without bias.

For CNN baseline models, the graph transformer module is removed and coordinates are directly projected form the 1D feature decoder output 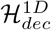.

#### VAE

To implement a VAE framework within our encoder-decoder architecture, we combines the encoder output by a horizontally average-pooling(HAP) the pairwise output feature and concatenate it with the scalar output before applying the mean and variance projection networks.

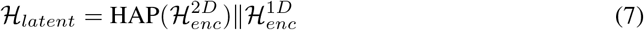

Where 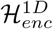 and 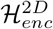 are the scalar and pairwise of the encoder output respectively and ∥ denotes concatenation. To produce the mean and variance of the latent variable we use two separate projections on *ℋ*_*latent*_.

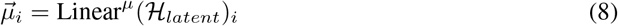

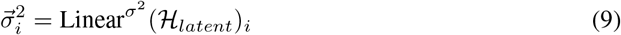

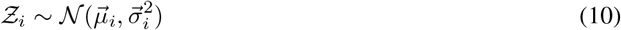

To produce the input scalar and pairwise features to the decoder networks, we take the outer product of the sampled latent representations.

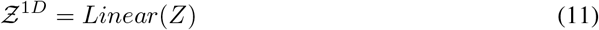

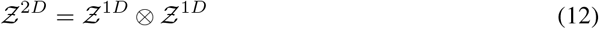

Where 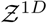 and 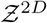 are the input to the scalar and pairwise feature decoder respectively.

#### Loss function and objectives

Let 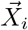 denote the backbone coordinates of residue *v*_*i*_. We predict backbone coordinates of 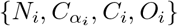 for each residue *i*. After predicting coordinates { *N*_*i*_, *C*_*αi*_, *C*_*i*_, *O*_*i*_ }, we follow (Jumper et al., 2021), and define the local Cα - frame of residue i as the rigid transformation 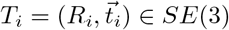 such that

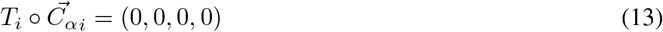

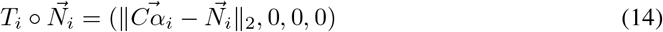

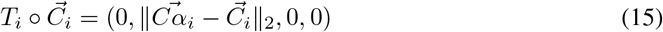

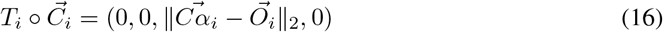

where 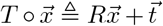. Assuming linear independence between the displacement vectors 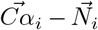 and 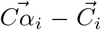, this transformation is unique and well defined. With a single local frame defined from each residue’s predicted coordinates, we are able to apply per-residue frame aligned point error (pFAPE) loss against the native coordinates and local frames as

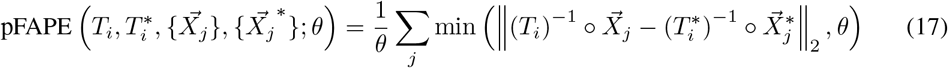

where *θ* = 10 is a threshold determining when the loss value should be clamped, and an asterisk is used to differentiate between native and predicted frames and coordinates. The pFAPE loss is averaged over all frames T_*i*_ and all atom types X to produce the final loss *ℒ*_*F AP E*_.

For the VAE loss, in addition to the FAPE reconstruction loss, we also compute the Kullback–Leibler divergence loss

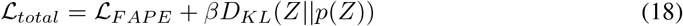

Where *D*_*KL*_ is the KL Divergence between the latent variable distribution and the prior multivariate Normal distribution.

#### Fixed backbone sequence design

We use the pre-trained fixed backbone design model of (McPartlon et al., 2022) to generate sequences from backbone structures. For each test target, we generate 500 conformational decoys and sequences are deranged for each of the decoys independently.

#### Structrue prediction oracles

We use the AF2 (Jumper et al., 2021) implemented by ColabFold (Mirdita et al., 2022) for structure prediction, for all structures, we use single sequence without MSA and set the number of recycles to three for all sequences. We make structure prediction to each sequence designed from the conformational decoys library and select sequence based on the prediction results.

### 5.3 Data

#### CATH4.2

We obtained the the CATH4.2 data from Ingraham et al. (2019) which contains 19,752 structures and structurally split with into train/validation/test sets by CATH fold annotations. To ensure sequence and structural independence between the train/test structures, the 40% non-redundant set is used and split is done on the topology/fold level.

#### Antibody Structures

For antibody structures, we used the structural antibody Database(SAbDab) obtained from Jin et al. (2021) which contains 1266, 1564, 2325 structures for CDR-H1, CDR-H2, and CDR-H3 respectively after filtering and splitting, there is no more than 40% sequence identity in the inpainted regions for each set of structures between the train/test structures. Please refer to Jin et al. (2021) for further details.

## Author Contributions

J.X. conceived and supervised the project and revised the manuscript. B.L. conceived the project, developed and tested the algorithm, analyzed and collected the results, and wrote the manuscript. M.M. assisted in writing the manuscript.

## Acknowledgments

We thank members of the Xu group for helpful discussions.

## A Appendix

### A.1 Additional Results

**Table 4:** Average structure similarity metrics of unmaksed structure reconstruction on models trained with different masking rate.

| Model | lddt↑ | TM↑ | RMSD↓ | Distance diff↓ | contact (S)↑ | contact(M)↑ | contact(L)↑ |
| --- | --- | --- | --- | --- | --- | --- | --- |
| CoordVAE(50% mask) | 0.809 | 0.918 | 1.370 | 0.584 | 0.830 | 0.790 | 0.762 |
| CoordVAE(60% mask) | 0.757 | 0.887 | 1.682 | 0.834 | 0.742 | 0.711 | 0.668 |
| CoordVAE(70% mask) | 0.753 | 0.883 | 1.700 | 0.994 | 0.714 | 0.667 | 0.670 |

**Table 5:** Average structure similarity metrics of linearly maksed input on models trained with different masking rate.

| Model | lddt↑ | TM↑ | RMSD↓ | Distance diff↓ | contact (S)↑ | contact(M)↑ | contact(L)↑ |
| --- | --- | --- | --- | --- | --- | --- | --- |
| CoordVAE (unmasked) | 0.639 | 0.711 | 5.705 | 2.662 | 0.306 | 0.290 | 0.304 |
| CoordVAE(50% mask) | 0.712 | 0.824 | 2.970 | 0.773 | 0.588 | 0.632 | 0.555 |
| CoordVAE(60% mask) | 0.672 | 0.792 | 3.287 | 1.070 | 0.512 | 0.531 | 0.479 |
| CoordVAE(70% mask) | 0.666 | 0.799 | 3.131 | 1.249 | 0.505 | 0.518 | 0.500 |

**Table 6:** Structure reconstruction performance of linearly masked input on model with different masking strategies.

| Masking rate = 50% |  | full |  | Masked region |  |
| --- | --- | --- | --- | --- | --- |
| Model | | lddt↑ | RMSD↓ | lddt↑ | RMSD↓ |
| CoordVAE (unmasked) |  | 0.597 | 5.682 | 0.363 | 19.056 |
| **CoordVAE(10AA linear)** |  | **0.714** | **1.738** | **0.527** | **3.729** |
| CoordVAE(spatial) |  | 0.705 | 1.809 | 0.551 | 3.689 |
| CoordVAE(Random) |  | 0.689 | 2.107 | 0.459 | 5.397 |
| CoordVAE(mixed mask) |  | 0.685 | 1.927 | 0.514 | 3.846 |

**Table 7:** Structure reconstruction performance of spatially masked input on model with different masking strategies.

| Masking rate = 50% |  | full |  | Masked region |  |
| --- | --- | --- | --- | --- | --- |
| Model | | lddt↑ | RMSD↓ | lddt↑ | RMSD↓ |
| CoordVAE (unmasked) |  | 0.391 | 8.820 | 0.261 | 15.386 |
| CoordVAE(10AA linear) |  | 0.585 | 3.208 | 0.417 | 5.689 |
| **CoordVAE(spatial)** |  | **0.670** | **2.098** | **0.537** | **3.606** |
| CoordVAE(Random) |  | 0.601 | 3.089 | 0.437 | 5.566 |
| CoordVAE(mixed mask) |  | 0.626 | 2.466 | 0.481 | 4.209 |

**Table 8:** Structure reconstruction performance of randomly masked input on model with different masking strategies.

| Masking rate = 50% |  | full |  | Masked region |  |
| --- | --- | --- | --- | --- | --- |
| Model | | lddt↑ | RMSD↓ | lddt↑ | RMSD↓ |
| CoordVAE (unmasked) |  | 0.308 | 9.910 | 0.283 | 12.149 |
| CoordVAE(10AA linear) |  | 0.359 | 7.114 | 0.307 | 9.870 |
| CoordVAE(spatial) |  | 0.561 | 3.124 | 0.497 | 4.834 |
| **CoordVAE(Ramdon)** |  | **0.743** | **1.461** | **0.658** | **2.348** |
| CoordVAE(mixed mask) |  | 0.632 | 2.319 | 0.540 | 3.715 |

**Figure 7:**
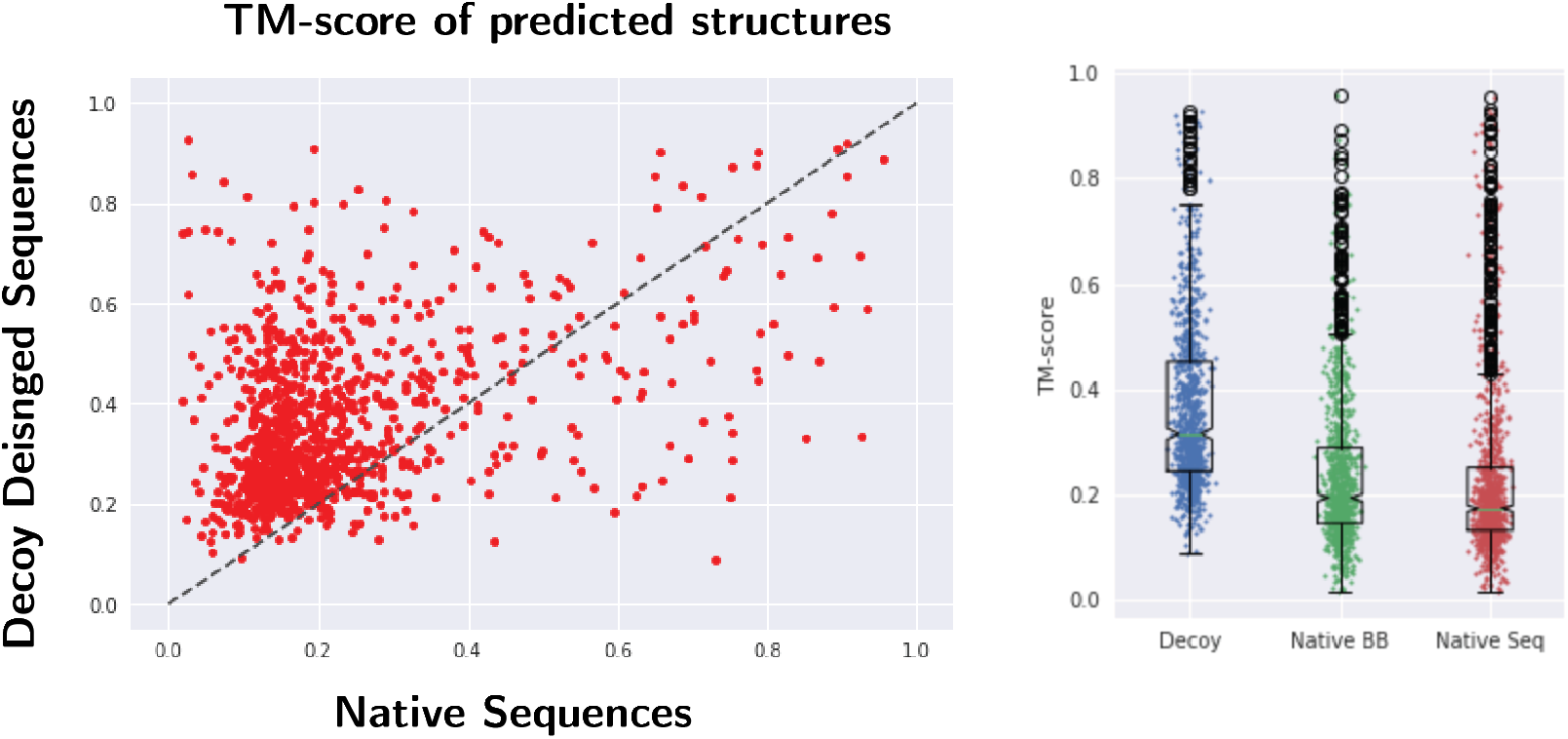
Scatter plots of the TM-score of the predicted structures from designed sequence and native sequence(Left). Distribution of TM-score across decoy designed sequences, fixed-backbone designed sequences and native sequences predicted by AF2.

**Figure 8:**
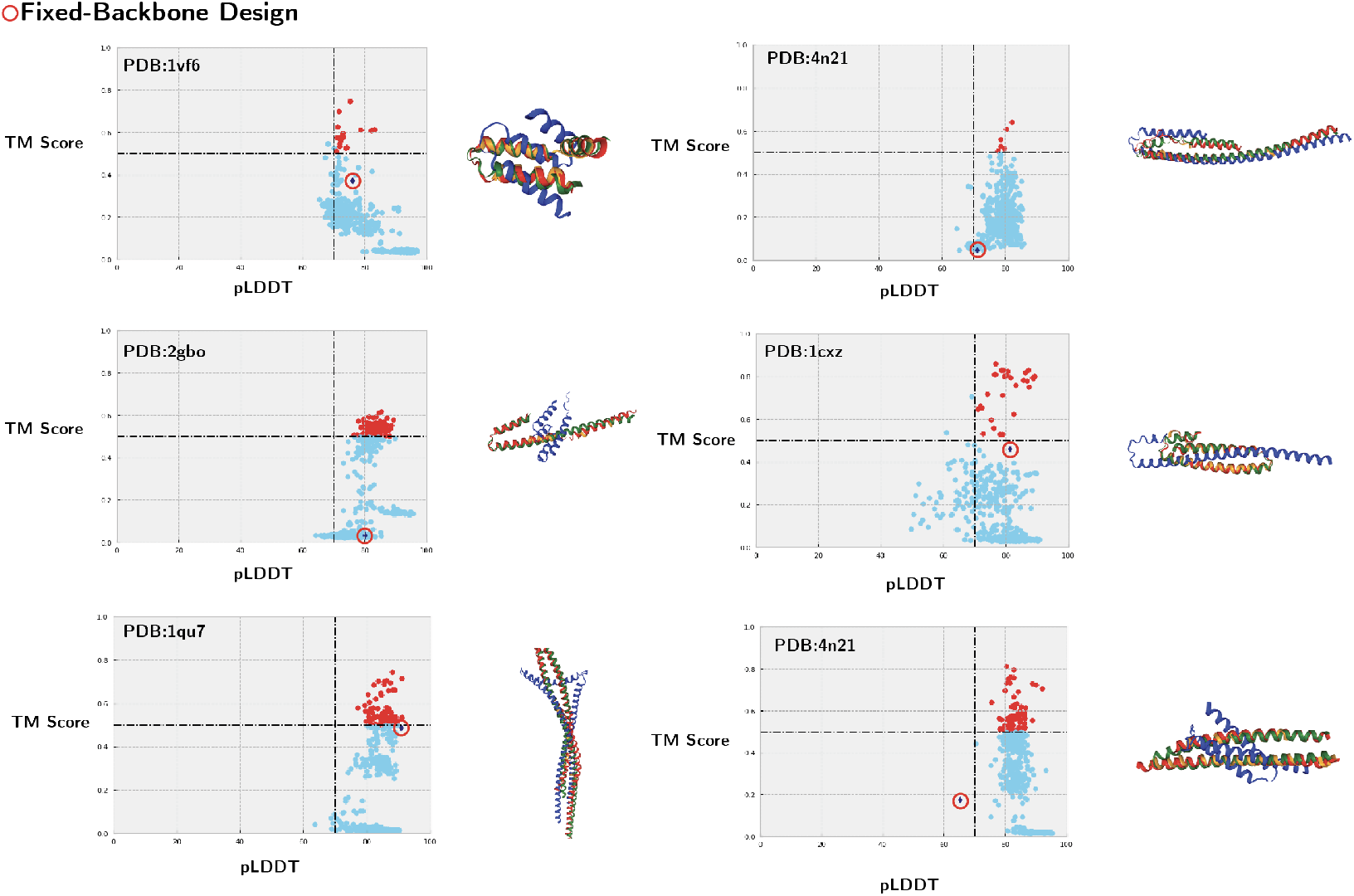
Example distributions of designed decoy sequence TM-score vs. pLDDT predicted by AF2. Fix-backbone designed sequence is circled in red circles. Successful designs are defined as pLDDT>70 and TM-score >0.5 which are colored red.

## Notes

### Competing Interest Statement

The authors have declared no competing interest.

